# Identifying branch-specific positive selection throughout the regulatory genome using an appropriate neutral proxy

**DOI:** 10.1101/722884

**Authors:** Alejandro Berrio, Ralph Haygood, Gregory A Wray

**Affiliations:** Duke University, 130 Science Dr, Durham, NC 27708, USA; Ronin Institute for Independent Scholarship, 127 Haddon Pl. Montclair, NJ 07043, USA

**Keywords:** Adaptation, Positive Selection, Analytical Method, *adaptiPhy*, proxy, neutral

## Abstract

Adaptive changes in *cis*-regulatory elements are an essential component of evolution by natural selection. Identifying adaptive and functional noncoding DNA elements throughout the genome is therefore crucial for understanding the relationship between phenotype and genotype. Here, we introduce a method we called *adaptyPhy*, which adds significant improvements to our earlier method that tests for branch-specific directional selection in noncoding sequences. The motivation for these improvements is to provide a more sensitive and better targeted characterization of directional selection and neutral evolution across the genome. We use ENCODE annotations to identify appropriate proxy neutral sequences and demonstrate that the conservativeness of the test can be modulated during the filtration of reference alignments. We apply the method to noncoding Human Accelerated Elements as well as open chromatin elements previously identified in 125 human tissues and cell lines to demonstrate its utility. We also simulate sequence alignments under different classes of evolution in order to validate the ability of *adaptiPhy* to distinguish positive selection from relaxation of constraint and neutral evolution. Finally, we evaluate the impact of query region length, proxy neutral sequence length, and branch count on test sensitivity.

## INTRODUCTION

An accurate and comprehensive characterization of the genomic distribution of adaptive substitutions is essential for understanding the genetic basis for trait divergence between species (Wayne and Simonsen 1998; Yang and Bielawski 2000; Nadeau and Jiggins 2010; Pardo-Diaz et al. 2015; Reilly and Noonan 2016). Tests for positive selection at the interspecies scale developed during the 1980s focused on ω, the ratio of nonsynonymous to synonymous substitution rates in protein coding regions (Li et al. 1985; Nei and Gojobori 1986). These methods were first applied at a whole genome scale soon after the release of reference genome assemblies for human, chimpanzee, and macaque (Lander et al. 2001; Clark et al. 2003; Iacobuzio-Donahue et al. 2003; Rhesus Macaque Genome Sequencing and Analysis Consortium et al. 2007), and provided the earliest relatively unbiased views of positive selection on protein-coding regions. At the same time, a growing appreciation for the contribution of regulatory mutations to adaptation (Hedrick and McDonald 1980; Prud’homme et al. 2006; Wray 2007) prompted the development of methods to test for positive selection in noncoding regions.

Two general approaches were devised to test for positive selection in the absence of a genetic code. One seeks regions that contain many substitutions along the human branch (since the most recent common ancestor with chimpanzees) but are otherwise highly conserved among mammals or vertebrates (Pollard, Salama, King, et al. 2006; Siepel et al. 2006; Bird et al. 2007; Bush and Lahn 2008; Prabhakar et al. 2008). The other seeks an elevated rate of substitution along the human lineage in a query region hypothesized to contain regulatory elements relative to a nearby reference (proxy neutral) region likely to contain few functional elements (Wong and Nielsen 2004; Haygood et al. 2007). Both approaches test for branch-specific accelerated substitution, but differ in the reference point against which they assess acceleration: the first tests for accelerated substitution within otherwise conserved regions against a putatively neutral region that is usually obtained from local Ancient Repeats (ARs) or fourfold degenerate sites (4D) (Pollard, Salama, King, et al. 2006; Bird et al. 2007; Bush and Lahn 2008; Prabhakar et al. 2008), while the second employs a putatively non-functional local intron of the genome as a neutral reference against which to identify branch-specific accelerated substitution (Wong and Nielsen 2004; Haygood et al. 2007). Wong & Nielsen (2004) defined the parameter ζ as the ratio of substitution rates in the query region to those in the associated neutral region; ζ is thus analogous to ω. To detect significance in the departures from neutrality, both approaches typically use maximum likelihood estimation and likelihood ratio tests (LRTs) that compare a null model allowing neutrality against an alternative model that additionally allows for positive selection.

These two general approaches have complementary strengths and weaknesses. The first approach is less sensitive, in that it doesn’t make use of an appropriate proxy for neutral regions along the human lineage. This approach allowed the discovery of Human Accelerated Regions (HARs) (Siepel et al. 2006). However, there is no reason to suppose adaptive evolution along the human lineage has been confined to regions under purifying selection in most or all other species, particularly since such regions constitute a small fraction of the genome (Figure 1A). Additionally, this method may fail to identify regions evolving under relaxation of constraint from those that have experienced positive selection. The first method has been implemented in *phyloP* (Hubisz et al. 2011), which is straightforward to execute and allows running selection tests using different approaches, such as LRT, SPH, Score and Genomic Evolutionary Rate Profiling (GERP) (Cooper et al. 2005; Rao 2005; Pollard, Salama, Lambert, et al. 2006; Siepel et al. 2006). *phyloP* has been extensively used for more than a decade, and has been applied to conserved DNA regions using neutral proxies based on four-fold degenerate (4D) sites (Pollard et al. 2010) or local ancient repeats (ARs) (Gittelman et al. 2015; Dong et al. 2016). The second method runs in *HyPhy* (Pond et al. 2005) and requires more analytical effort than *phyloP*. On the other hand, it is more broadly applicable because it can be applied to any genomic region regardless of whether that region was previously under functional constraint.

**Figure 1.**
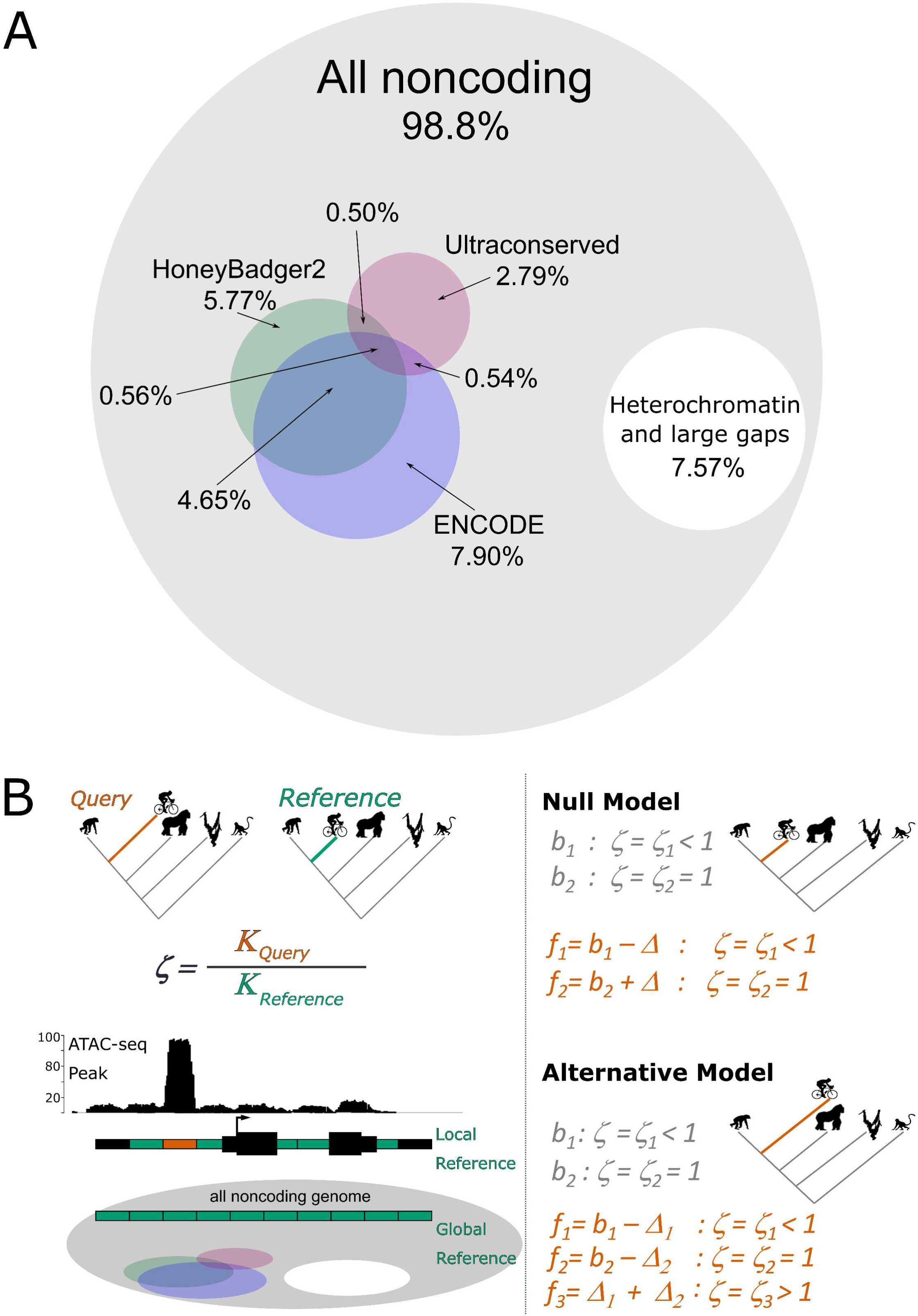
Overlaps between functional regions of the human genome and the evolutionary model. **A.** *Left*, Venn diagram showing percentage of the human genome that overlaps with known non-coding functional annotations and conserved regions of the human genome. *Right*, scaled Venn diagram showing the proportion of conserved regions with respect to known functional annotations and gapped DNA representing telomeric and centromeric sequences (white). **B.** Graphic summary of our improved method. *Left panel*, the evolutionary ratio “ζ” is computed as the ratio of the substitution rate in a query (*K*_query_) with respect to the substitution rate in a reference (*K*_reference_) region. Queries can be obtained from functional annotations such as ATAC-seq or ChIP-seq peaks (red box), while reference alignments can either be taken by sampling local non-functional elements in the vicinity of the query or from a genome-wide random sampling of non-functional and putatively neutral regions of the genome (green boxes). *Right panel*, to obtain a *P*-value describing the significance of a given evolutionary ratio ζ, we compare the ‘null’ model against an “alternative’ model on a foreground branch (red). Where a proportion of sites in the query region (f_3_) evolve under positive selection, while the background, they evolve by purifying selection or putatively neutral in both query and reference; *b* refers to the background and *f* to foreground branches (modified from Haygood et al. 2007). Δ refers to the estimated proportion of DHSs with negative selection on background lineages and neutral on foreground lineage in the null model. Δ_1_ refers to the estimated proportion of DHSs with negative selection on background lineages and positive selection on foreground lineage, and Δ_2_ refers to estimated proportion of DHSs that are neutral on background lineages and positive selection on foreground lineage in the alternative model.

At the time these approaches were first developed, very little information existed about the location of functional elements in any genome. This limited the ability to identify suitable proxy neutral regions, i.e., those likely to be free from either purifying or positive selection. Inadvertently using constrained or accelerated regions as neutral proxies can potentially introduce artificial adaptive signals or reduce sensitivity, respectively. In addition, not knowing the location of regulatory elements meant that testing for positive selection at a genome-wide scale was intractable due to the need for massive correction for multiple testing. Prior to the invention of functional genomic assays for chromatin status, the best method for identifying putative regulatory elements was sequence conservation (Pollard et al. 2006).

The ENCODE project (Myers et al. 2011; Bernstein et al. 2012) and other efforts (Kundaje et al. 2015) to identify regulatory elements throughout the human genome mean that it is now possible to focus tests for positive selection on likely functional noncoding elements and to identify appropriate proxy neutral regions. Highly conserved noncoding regions overlap with only about 1.09% of the ~3 million known DNase I Hypersensitive Sites (DHSs) in the human genome (Thurman et al. 2012) and just 1.06% of ~1.7 million enhancers and promoters from the HoneyBadger2-intersect regions published by the ENCODE and Roadmap Epigenomics projects (Figure 1A). Accordingly, it seems likely that a substantial fraction of functional DNA elements do not occur in regions within regions of strong conservation. At the same time, ~18.3% of the human genome currently has no known regulatory and functional annotation, despite extensive study, providing a principled basis for choosing proxy neutral sequences that may be superior to 4D sites and ARs.

Here we introduce *adaptiPhy*, an improved analytical method that can test for branch-specific directional selection on any collection of query segments based on accurate alignments from three or more species. We implemented a series of technical and computational modifications to our previously published method (Haygood et al. 2007) using openly available software from PHAST (Siepel et al. 2005; Hubisz et al. 2011) and functional genomic datasets from ENCODE (Rosenbloom et al. 2011). We tested the performance of *adaptiPhy* in regions that have been already tested (i.e., published Human Accelerated Regions, or HARs) and among simulated sequences evolving at neutral rates, positive selection in one branch of the tree, and relaxation of constraint. Significant improvements include: 1) better genome-wide representation of putatively neutral proxies based on functional annotations; 2) ability to test for selection in nearly any noncoding region of the genome, including functionally dense regions; 3) the ability to increase stringency by filtering reference sequences; and 4) improved understanding of how branch number and the size of query and reference regions impact test sensitivity. We demonstrate that this approach can be applied productively to focal collections of genomic regions commonly encountered in contemporary genomics and genetics research, such as the open chromatin landscape of a specific cell type or trait-associated regions from a GWA study.

## RESULTS

### Global nonfunctional sequences provide appropriate neutral reference sequences

To identify appropriate proxy neutral regions, we began by identifying all non-functional regions (NFRs; see Methods for inclusion criteria) of length 300 bp (similar in length to many regulatory elements) throughout the human genome. We then tallied the number of NFRs located within 10, 40 and 100 kb of a set of 1000 random DHS sites, non-coding Human Accelerated Elements (ncHAE), and a control subset of “global” NFRs from throughout the genome. On average, there are only 7.8 local NFRs per DHS, 28.8 NFRs per ncHAE, and 64.4 NFRs per global NFR (Figure 2A). Moreover, 58.5 % of DHSs and 43.3% of ncHAEs had *no* local NFRs within 100 kb. Thus, the number of local NFRs that can be used as neutral reference regions is often insufficient for extensive testing of positive selection. Some previous studies used ancient repeats (ARs) as neutral proxies (e.g., Dong et al. 2016). We found that on average, there are only 3.3 ARs within 100 kb per DHS, and 44% of DHS regions had no AR within 100 kb. Thus, identifying sufficient ARs to use as a local reference for each DHS is also difficult.

**Figure 2.**
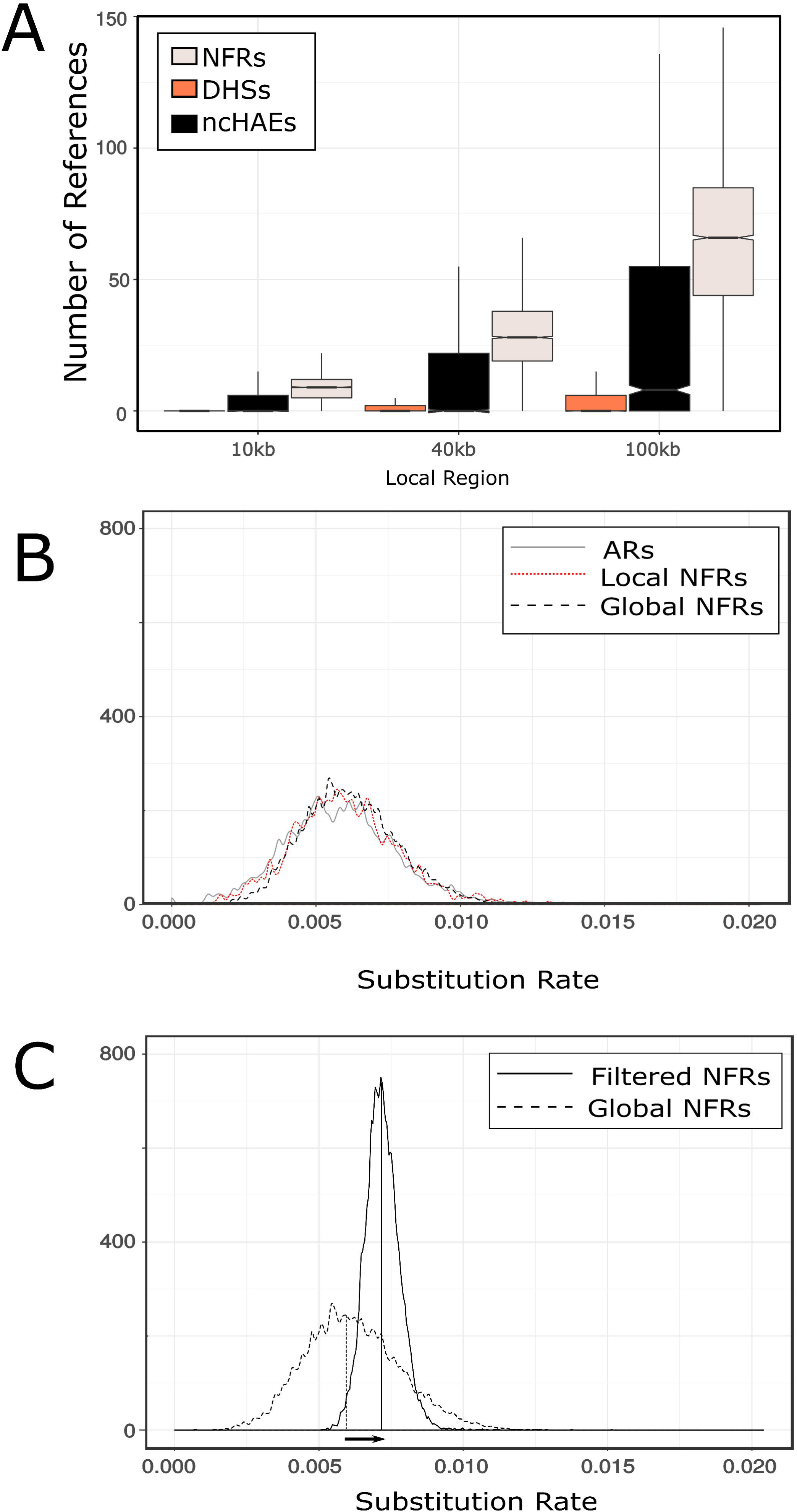
Finding a neutral proxy. **A.** Distribution of the number of local reference alignments around each DNA element within three different distances: 10kb, 40kb and 100 kb. **B.** Density distribution of relative substitution rates among concatenated Ancestral Repeats (ARs) around 100 KB of each DHS element in our list, concatenated local NFRs around each DHS, concatenated global NFRs before filtering out trees with low and high substitution rates. **C.** Density distribution of relative substitution rates in the concatenated global NFRs before and after filtration step. The arrow depicts the change in the median distribution of substitution rate of global reference alignments before to after filtering.

Next, we asked whether local ARs, local NFRs, or global NFRs elements can be used to build an appropriate reference for testing positive selection. To build a local reference region, we concatenated all the NFRs or ARs within 100 kb of a given query region. Next, we computed the substitution rate of each concatenated sequence of local ARs, local NFRs, and NFRs across the genome. For a set of query regions in our sample, we found a wide distribution of substitution rates among concatenated local references including local NFRs, global NFRs, and ARs (Figure 2B).

We thus sought to test whether filtering global NFRs by their relative substitution rate respect to the entire tree can provide an improved neutral proxy for estimating positive selection. We filtered out global NFRs representing the top and bottom quartiles of relative substitution rates (Figure S1), then concatenated 10 NFRs per query. When we compared the substitution rates of this new set of putative neutral references, we found that the distribution of substitution rates of the filtered global NFRs is narrower and its median is skewed to the right (Figure 2C). This suggests that global NFRs can provide a set of putatively neutral elements if appropriately filtered. This approach also allows conservativeness and sensitivity to be modulated by tuning the filtration step accordingly. Moreover, given that we use relative values of substitution rate, this filtering step can applied to any region of the genome regardless of the amount of functional annotation.

To validate the effect of local and global NFRs when testing for positive selection, we sampled three sets of queries: (1) widespread and specific DHSs (open in >124 and exactly 1 ENCODE cell types, respectively); (2) a set of ncHAEs to be used as positive controls; and (3) a set of putatively non-functional DNA elements to be used as negative controls. The correlation in *P*-values is high among the 3,531 DHSs that could be compared using both local and global neutral proxies (Spearman’s Rank test rho = 0.80; *P* < 2.2×10^−16^; Figure 3A). Of these, only 2.63% scored high for positive selection (*P* < 0.05) for global proxies, while 5.12% scored high for positive using the local proxy alone. Likewise, the correlation of *P*-values in the global and local sets is high among the 1,291 ncHAEs that could be tested using both local and global proxies (Spearman’s Rank test rho = 0.86; *P* < 2.2×10^−16^). Of these, only 25.33% of the ncHAE regions tested positive globally, while 39.04% tested positive for selection using the local tests (Figure 3B). Thus, local proxies in general identify more putative cases of positive selection but have limited applicability within function-dense regions of the genome while global proxies can be used to test any query region but possibly with lower sensitivity.

**Figure 3.**
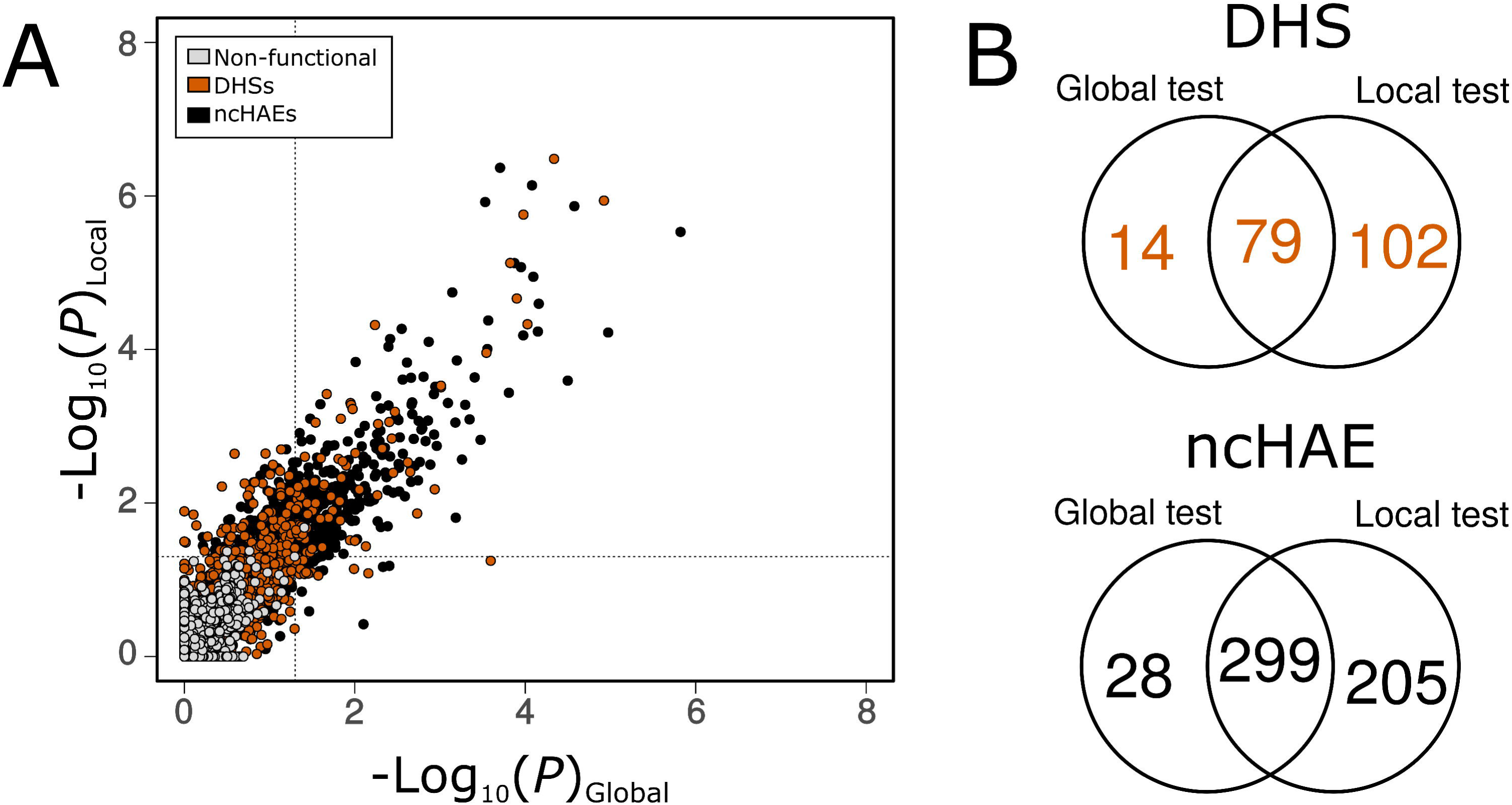
Global proxy as a useful neutral proxy. **A.** Correlation between local and global tests of selection among different classes of DNA elements; the Spearman rank correlation coefficient is highly significant (*P* < 2.2 x10^−16^) and very high (rho > 0.80) for DHSs and ncHAEs (rho > 0.86). Inner dashed lines depict a significance level of *P*=0.05. **B.** Venn Diagram of the overlap of regions scoring high for positive selection using the global test vs the local test for DHS elements (top) and ncHAEs (bottom).

### Sensitivity is high given practical query length, reference length, and branch number

Earlier, we observed that the distribution of substitution rates among all global reference regions used in this study is narrow and high (Figure 2B). More specifically, we observed an average human branch length of 0.0072 substitutions per site using global neutral proxies, which is appreciably faster than the average substitution rates of local elements (average branch length = 0.0055). We evaluated the effect of query and neutral proxy length in the estimation of positive selection, using both real and simulated data. To evaluate the effect of reference length on sensitivity, we tested reference alignments of 300 bp, 900 bp, 3 kb, 9 kb, and 30 kb (Figure 4A). As expected, the median of the branch lengths of the global references does not increase or decrease as they get longer; rather, they reach an equilibrium at 0.00725 with reduced variation (Figure S5).

**Figure 4:**
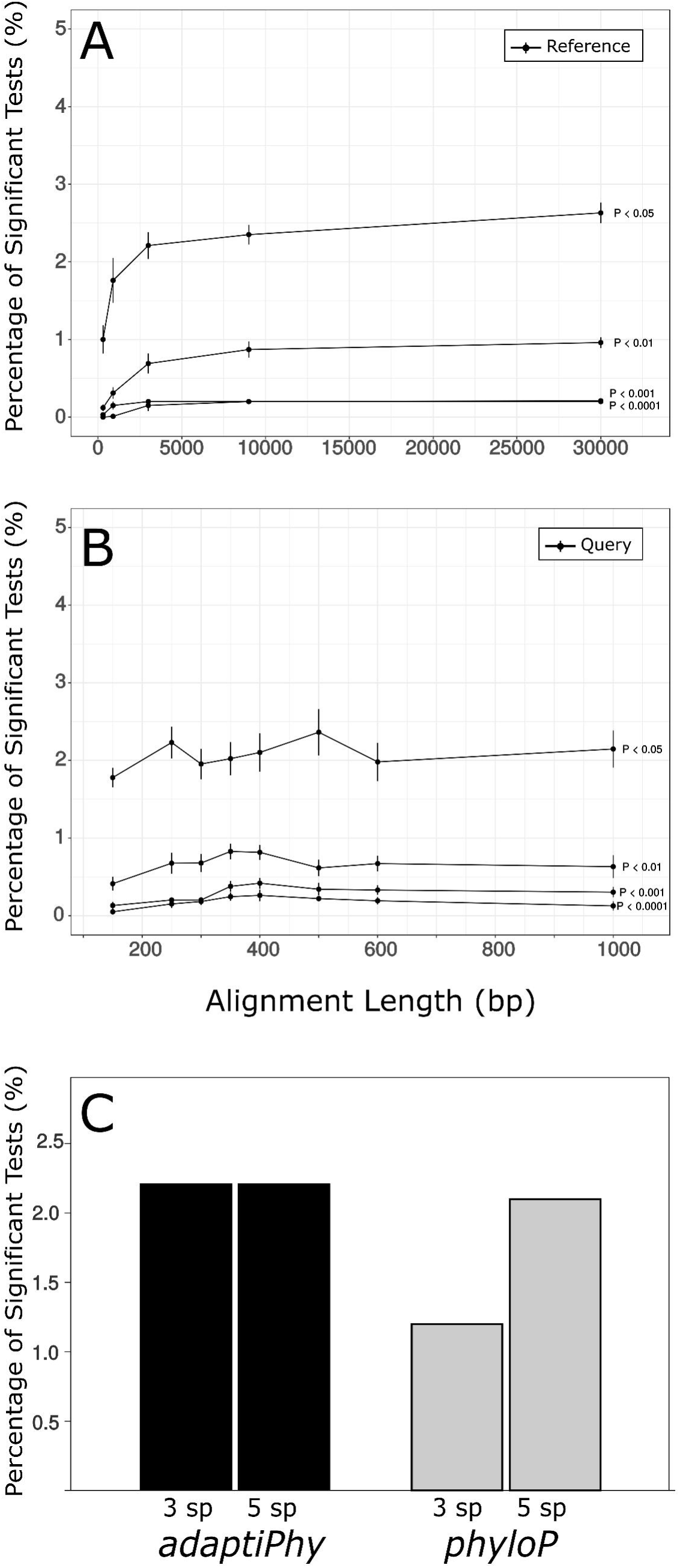
Sensitivity test and the effect of reference and query length, and number of species in the estimation of selection in a subset of 1000 random DHSs. **A.** Effect of query length on the power to identify selection at different significance levels. **B.** Effect of reference length on the power to identify selection at different significance levels. **C.** Percentage of significant tests in a sample of 1000 DHSs that were in *phyloP* and *adaptiPhy*. Bars labeled as 5 sp depict tests done for five branches, while 3 sp labels depict three-species tests.

Functional genomic approaches such as ChIP-seq and ATAC-seq identify putative regulatory regions with window lengths that are usually between 150-800 bp and skewed towards shorter lengths (Crawford 2005; Birney et al. 2007; Song and Crawford 2010; Shibata et al. 2012; Thurman et al. 2012; Bryois et al. 2017). In order to assess the ability of *adaptiPhy* to identify positive selection throughout the biologically meaningful range of putative relative element sizes, we tested the effect of query lengths. Importantly, the ability to detect selection is not strongly affected by differences in query length, and remains similar down to ~150 bp (Figure 4B). This finding suggests that our set of reference sequences is able to detect signatures of positive selection in regions where the query has been predicted to be longer than the actual functional element under selection, and in particular across most of the size range of known regulatory elements and open chromatin regions in the human genome.

Finally, we tested the impact of using three or five species to detect positive selection, as more branches might be expected to provide a better estimation of the background substitution rate in the reference. We found that adding one or two species above the minimum of three (two ingroup and one outgroup) provides only a negligible improvement in sensitivity of *adaptiPhy* (Figure 4C). In contrast, the sensitivity of *phyloP* is more dependent on the number of species, improving markedly with additional taxa (Figure 4C). Thus, *adaptiPhy* may be preferable in situations where the minimum number of reference genome assemblies is available.

### The test discriminates between three different types of selection

To determine whether *adaptiPhy* can correctly detect selection under different evolutionary scenarios, we simulated both query and reference alignments undergoing different types of evolution, including neutral in both the background (BG) and the foreground (FG), constraint in the BG but neutral in the FG (i.e., relaxation of constraint), and neutral in the BG but positive selection in the FG. We found that *adaptiPhy* accurately discriminates positive selection from relaxation of constraint and neutral evolution (Figure 5). Of the simulated neutral regions, 2.3% were incorrectly identified as positive selection using *adaptiPhy* compared to 4.1% using *phyloP*. In addition, *phyloP* fails to distinguish positive selection from relaxation of constraint in almost half of the simulated cases, while our approach fails in only 2.7% of cases. In contrast, of the simulated regions under positive selection, *adaptiPhy* and *phyloP* identified 99.6% and 98.8% of sequences simulated to be under positive selection, respectively (Figure 5). Moreover, the sensitivity of *adaptiPhy* to detect positive selection is 1 and specificity is 0.97, while the sensitivity of *phyloP* is 0.99 and specificity is 0.73. While it is difficult to evaluate whether the same would occur with real data, these results from simulations suggest that *adaptiPhy* may produce a lower false positive rate than *phyloP*, particularly for instances of relaxed selection.

**Figure 5:**
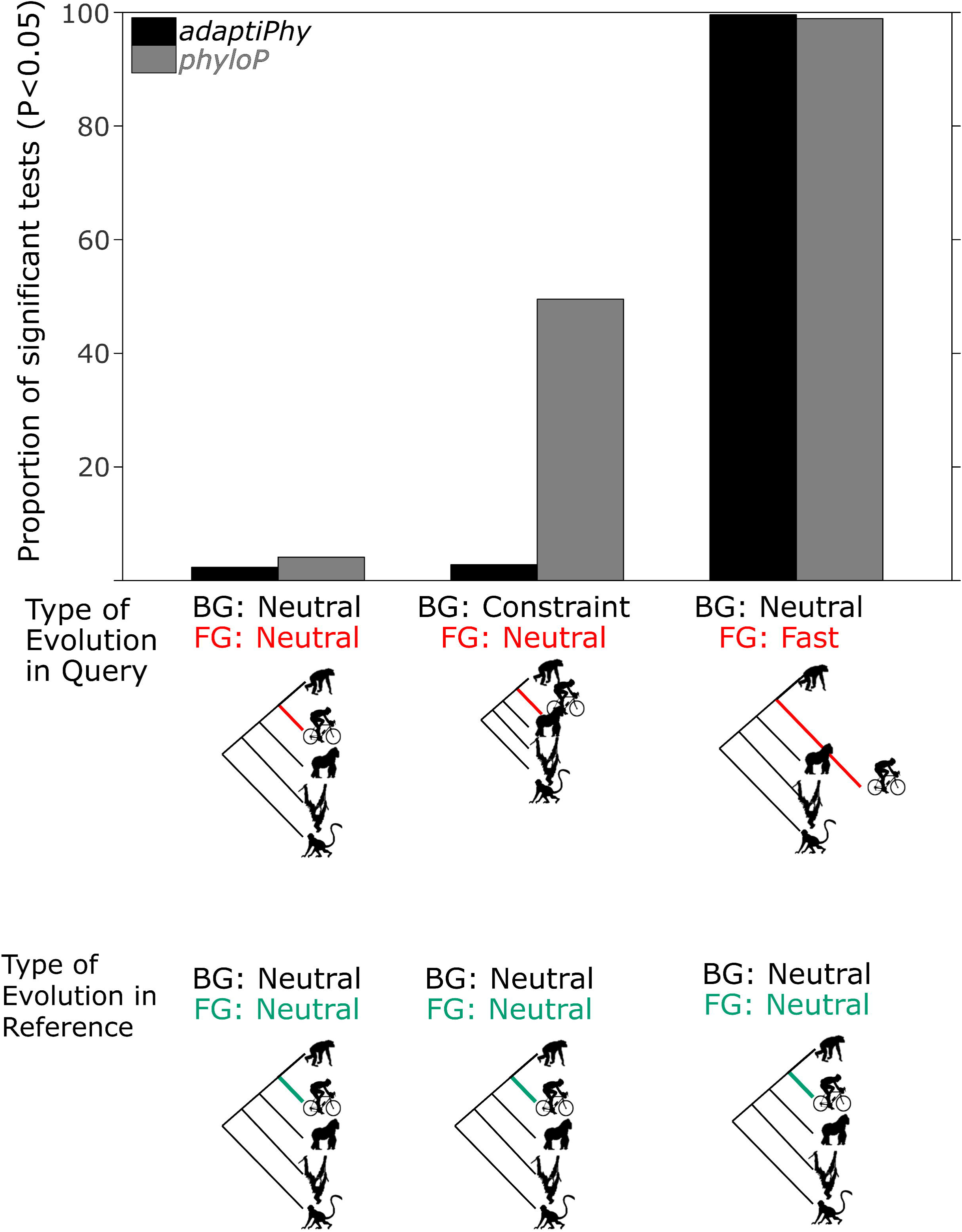
Specificity test of *adaptiPhy* and *phyloP* for three types of evolution. All trees and alignments were simulated to be evolving by neutral evolution (left), relaxation of the constraint (center), and positive selection (right). The left and center simulations give an idea of the amount of type I, while the right simulation provides a estimation of the type II errors.

### The test reconfirms many previously identified Human Accelerated Elements

To test whether *adaptiPhy* can replicate results from previous scans for positive selection in noncoding regions during human origins (Pollard, Salama, King, et al. 2006; Bird et al. 2007; Bush and Lahn 2008; Prabhakar et al. 2008), we queried a consolidated set of 2,649 noncoding Human Accelerated Elements (ncHAEs) (Capra et al. 2013). Since our findings indicate that our test does not dilute signals of positive selection when the query regions is between 150 bp and 1 kb, and given that most ncHAEs are relatively short (67% are shorter than 300 bp), we normalized query length by capturing sequence up to 300 bp centered on each ncHAE. At a significance level of 0.05, we confirmed 26.44% of previously reported ncHAEs. More broadly, the distribution of *P*-values for previously reported ncHAEs is skewed toward 0; the peak at the lower end of the distribution is pronounced (Figure S4C), as expected, this distribution is more skewed towards 0 using *phyloP* using our set of neutral references (Figure S4D). Indeed, these results are consistent with acceleration in the human lineage when compared with testing for positive selection on the chimpanzee branch.

Interestingly, there is a substantially higher degree of overlap between the regions we replicated in the consolidated set of ncHAEs and those in each of the published studies than between any two of them (Figure S6). For instance, we validated nearly 55% of the ncHAEs identified by Bush and Lahn (Bush and Lahn 2008), a significantly higher fraction than any of the other studies were able to validate (Fisher’s Exact Test, Two-sided, *P* = 0.0007). Of the other methods that were used to identify human accelerated elements, Bush and Lahn’s is the most methodologically similar to ours, although they used ancient repeats within 750 kb surrounding each ncHAE as the neutral proxy. Only four loci were identified as ncHAEs in all four prior studies, and all four were replicated here (Table S1 and Figure S7). One of these is *BNC2*, a gene that may be responsible for skin pigmentation differences between humans and other primates (Sankararaman et al. 2014; Vernot and Akey 2014).

We also scanned a region of 100 kb containing one of the genes that was significant for positive selection in the published studies (Pollard, Salama, King, et al. 2006; Bird et al. 2007; Bush and Lahn 2008; Prabhakar et al. 2008) and the present study using a sliding window. We found additional signals of positive selection around the *NEIL3* locus that were as strong as the single ncHAE element that was originally identified (Figure 6). The striking clustering of signals of positive selection around this locus suggest that transcriptional regulation of *NEIL3* changed extensively during human origins and highlights the ability of *adaptiPhy* to identify signatures of positive selection that other methods may miss.

**Figure 6:**
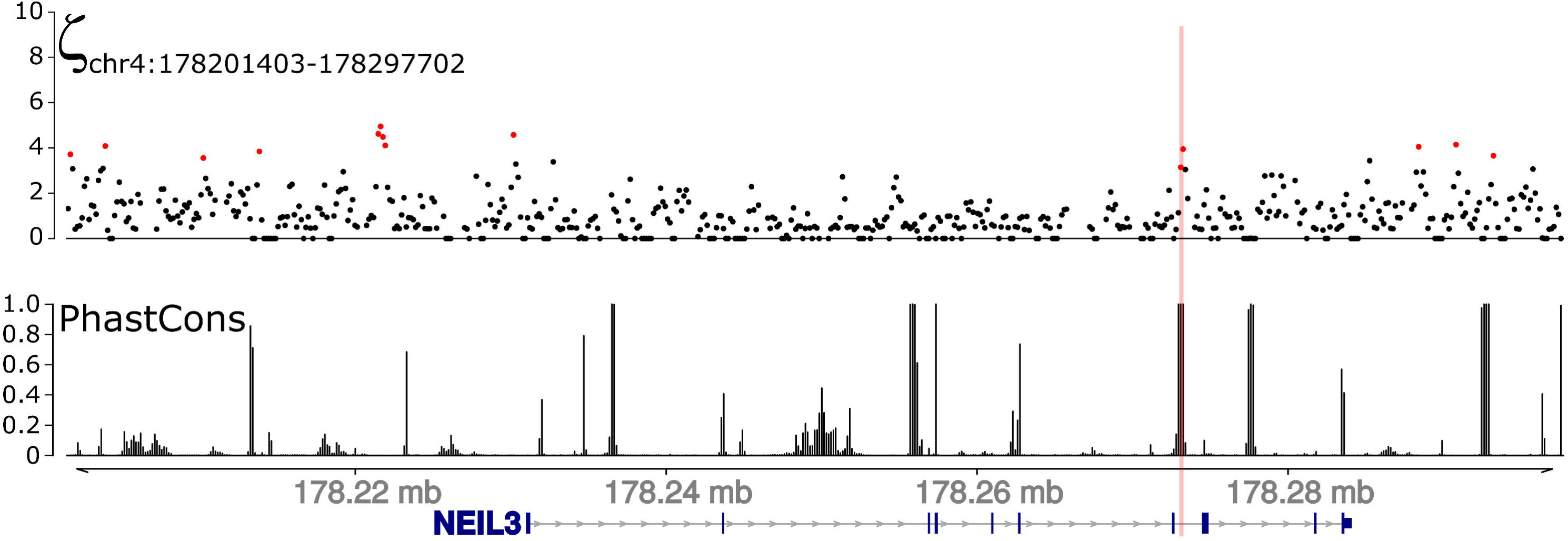
Sliding windows analysis of the rate of evolution along a genomic region containing a ncHAE. Distribution of positive selection as ζ (top panel), and vertebrate conservation scores (bottom panel) along a 100 kb region surrounding the gene NEIL3. This locus is located nearby one of the human accelerated elements identified in our test and across all four of the cited studies (Figure S7). The location of this ncHAE is highlighted in pink, and the red dots represent windows of 300 bp were ζ scored significant for positive selection (P < 0.05).

## DISCUSSION

Rigorous testing for selection depends upon accurate identification of reference regions that represent neutral rates of evolution. Prior studies have used four-fold degenerate (4D) sites from coding sequences (Pollard, Salama, King, et al. 2006), surrounding non-coding sequences (Gittelman et al. 2015), concatenated conserved LINE elements (ancient repeats; ARs) (Dong et al. 2016), and non-first intron sequences (Haygood et al. 2007) as neutral references. However, regulatory elements are often located in regions that are dense with functional annotations and thus potentially constrained or near sites evolving under weak selection (Figure 2A). Consequently, identifying sufficient non-functional regions to use as a neutral proxy in the vicinity of most open chromatin sites is simply not possible in primate genomes and the problem is even more acute in organisms with higher gene density. To address this limitation, we used the dense functional annotation of the human genome to identify a set of putatively non-functional regions that can be used as an appropriate neutral proxy.

We then filtered this set of putatively non-functional regions to address the challenge of rate heterogeneity. For the entire set of DHS elements in our sample, we found a wide distribution of substitution rates among concatenated local samples including local NFRs, raw global NFRs and ARs (Figure 2B). This means that there are many such regions in which the substitution rate on the human branch is substantially slower or faster than average and thus potentially confounding when used to test for positive selection. These observations reflect generally less availability of non-functional elements in regions where DHSs and ncHAEs are more common and slower rates of evolution nearby each functional element, but also the possibility that some ancient LINE elements in the vicinity of DHSs are under functional constraint or evolving under high mutation rates. Indeed, many studies have shown that ARs can become co-opted as local regulatory elements (Greene et al. 2005; Kamal et al. 2006; Maumus and Quesneville 2014; Zeng et al. 2018). Consistent with this interpretation, we found that the proportion of reference ARs with at least one multitissue eQTL or brain specific eQTL is almost three times as large as in a set of global non-functional sequences (Figure S3). The relative absence of local non-functional elements may also increase the effect of incomplete lineage sorting in the reference sequence, producing an increase of false positives and negatives for query elements if the reference foreground sequence is evolving at slower or faster rates than the background.

Together, these results suggest that using local non-functional elements and ARs can both contribute to misestimation of selection. At the same time, ARs have the tendency to overestimate the rates of evolution of query alignments compared to a set of neutral references built from global NFRs (Figure S2). Our findings also suggest that the local tests have a tendency to overestimate positive selection, probably because they contain unknown functional sites and thus underestimate the neutral rate. As expected, most putative non-functional elements appear neutral in tests for selection, with only one testing positive for selection using both local and global references. Overall, these results suggest that global proxies are capable of testing more genomic regions, with an important gain in accuracy as we sample neutral regions based on their distribution pattern relative to the tree rates. As a consequence, every query will be tested against a reference region in which the foreground rates are not evolving faster or slower than the background branches of the tree. Local proxies can be more sensitive by increasing the number of sites that are under positive selection, some of which may be falsely called by introducing reference sequences that are in constraint and are thus evolving at slower than neutral rates. Since incomplete lineage sorting can be a potential confounding factor for both local and global neutral proxies, our filtering of global sequences based on relative branch lengths and concatenation of non-functional elements provides a useful strategy to control this issue (Tonini et al. 2015).

We also observed that among DHSs and nonfunctional DNA elements, the distribution of *P*-values tends towards the upper limit (*P*=1). This means that most open-chromatin regions are probably evolving neutrally or under purifying selection because the maximum likelihood estimate of the null and the alternative are exactly the same when *P*=1 (Figure S4 C). In contrast, most tests on ncHAEs show a distribution of *P*-values strongly skewed towards zero with a small but important density peak at 1, suggesting that at least 92 out 2,416 regions originally identified as ncHAEs appear to be evolving neutrally or by purifying selection in the human branch when using global proxy. Some of these may reflect errors in the early reference assemblies that were used in prior studies. And indeed, our results are generally consistent with their findings (Pollard, Salama, King, et al. 2006; Bird et al. 2007; Bush and Lahn 2008; Prabhakar et al. 2008), identifying many of the same regions as being under positive selection. Moreover, our results suggest that the distribution of *P*-value is strongly dependent on the genomic partition: DHSs and our set of non-functional elements scored nearly 2.5% and 0.02% of sites under positive selection respectively, confirming the expectation that DHSs in general are more often subject to positive selection than nonfunctional DNA elements. This fraction is higher than previously reported (Gittelman et al. 2015; Dong et al. 2016). The likely reason is that these prior studies underestimated the proportion of sites under selection because the AR elements used as neutral proxies could be evolving under higher substitution rates than the rest of the genome (Figure S2).

By testing reference alignments of different lengths, we show a diminishing return as these proxies get longer. We recommend using reference alignments between 3 and 9 kb (Figure 4A), as longer alignments require more computing power while providing only minimal additional sensitivity. Reference alignments shorter than 3 kb introduce more variation in estimation of ζ and thus increase the risk of false positives and negatives. We also tested the effect of query region length and found little difference in sensitivity between regions of 150 bp and 1000 bp, which means our approach can be applied to the vast majority of putative regulatory elements. We recommend caution in extending query region length indefinitely because this risks combining multiple functional elements with distinct rates of substitution.

Because differences in the rate of substitution in the reference has a strong influence on sensitivity, using global references can be valuable, particularly in regions of the genome that are densely populated with genes or other functional elements. In addition, a strong correlation exists between the distributions of *P*-values found in our test of DHSs (Spearman’s Rank test rho = 0.78; *P* < 2.2×10^−16^) and ncHAEs (Spearman’s Rank test rho = 0.86; *P* < 2.2×10^−16^), and their corresponding rates of evolution (ζ) (Figure S4). This result suggests that both *P-*value and ζ should be used together to score, describe and visualize regions under positive selection (Figure 6). Here, we used the conventional *P* < 0.05 cutoff to nominate query regions likely under positive selection. However, more stringent nominations are possible using lower cutoffs, such as False Discovery Rate (Storey and Tibshirani 2003).

Our framework can be used to detect sequence outliers under a high substitution regime while controlling for relaxation of constraint. We propose that most signals that are detected reflect true instances of positive selection. However, locally elevated mutation rate is a potential confound. To investigate whether local differences in mutation might produce a false positive, we recommend investigating the amount of common and rare variation present in a query region by consulting dbSNP or 1000 genomes databases. Rare variants are often used as a proxy for both mutation rate and recombination rate (Nei 1977; Slatkin 1985; Nachman 2001; Nachman 2002), making this a straightforward way to flag queries that score high for positive selection but are under a higher or lower mutational regime. Moreover, finding an excess of common variants under high-to-intermediate frequencies in a given region that scores high for positive selection is potentially a useful way to identify previously unknown regions under persistent balancing selection. Indeed, we found that the most variable element under positive selection is located in the MHC region (Supplementary Data), a region known to be under balancing selection (Takahata et al. 1992).

In this study, we used the human (hg19), chimpanzee (panTro4), gorilla (gorGor3), orangutan (ponAbe2), and macaque (rheMac) reference assemblies. It is highly likely that none of these reflect the ancestral state for their respective species. Depending of the number of individual genomes that were assembled into the reference, some regions may contain concentrations of rare or common derived alleles. This will artificially increase the apparent interspecies substitution rate. To control for this potential confound, we suggest including both common and rare variation that is known for the branch of interest and build local alignments with reference genomes that have used ancestral state data to correct for intraspecific variation.

Together, the improvements introduced here improve the ability to identify proxy neutral regions and increase the ability to modulate the sensitivity and conservativeness of branch-specific tests for positive selection in noncoding regions. The test is sensitive across nearly the entire range of annotated functional regulatory elements, dropping only for elements <150 bp in length. It is possible to apply these tests to nearly any noncoding region of the genome, even those in functionally dense locations.

## MATERIALS AND METHODS

Testing for branch-specific positive selection using our approach requires at least one reference alignment in which all branches of the tree are evolving at putatively neutral rates, and one query alignment, which can be obtained from any genomic region of interest, such as putative open chromatin regions or segments of a GWAS peak (Figure 1B). First, we downloaded the 100-way multiple alignment from the University of California Santa Cruz (UCSC) website (http://hgdownload.soe.ucsc.edu/goldenPath/hg19/multiz100way/) and several annotations of functional DNA elements from the ENCODE Project at UCSC, including: 5’ and 3’ UTRs, total human mRNA, lincRNAs, microRNAs, sncRNAs, short repeats, CpG islands, etc. We also enriched this list of functional elements with a set of HoneyBadger2-intersect promoters and enhancers from reg2map annotated by the Broad Institute and Epigenomics Roadmap project. Then, using *maf_parse* implemented in PHAST (Hubisz et al. 2011), we transformed the 100-way alignment into a smaller 5-way genome-wide alignment in MAF format that included only our focal species human (hg19), chimpanzee (panTro4), gorilla (gorGor3), orangutan (ponAbe2), and rhesus macaque (rheMac3). To draw reference alignments and non-functional regions, we generated a masked 5-way MAF alignment with a BED file containing all known functional DNA regions using *maf_parse* with the optional command *--mask-features*. To mask the genome, we used a merged BED file that included 5’ and 3’ exons, all coding and non-coding RNAs, vista enhancers, roadmap and ENCODE regulatory elements and promoters, CpG repeats, microsatellite sequences and simple repeats. Interestingly, the remaining non-functional fraction of the genome covered only ~20.5% of the genome. To draw query alignments, we used used *msa_split* (Hubisz et al. 2011) to draw alignments from non-coding human accelerated elements, ncHAEs (Pollard, Salama, King, et al. 2006; Bird et al. 2007; Bush and Lahn 2008; Prabhakar et al. 2008; Franchini and Pollard 2017), a random subset of non-functional regions (as defined below), and DNA Hypersensitive Sites (DHSs) from 125 human cell types and tissues (Thurman et al. 2012).

To select non-functional regions (NFRs) for our analyses of positive selection, we randomly chose around two million (1,893,795) non-overlapping segments of 300 bp or longer from the non-functional fraction of the genome described above, collectively amounting to 18.3% of the genome. Subsequently, we excluded all the alignments containing any masked regions with functional sequences, CpG islands, repetitive elements, assembly gaps and heterochromatin sequences (masked as Ns), and/or missing sequences (masked as asterisks or dashes.) Next, we computed branch lengths of each of the tree branches using the tool *phyloFit* available in PHAST (Hubisz et al. 2011), *phyloFit* computes a tree with branch lengths and a substitution rate matrix by fitting a tree model into a multiple sequence alignment using maximum likelihood. Within the remaining sample of 92,160 alignments from the last filtration step, we excluded any alignments in which the human branch was evolving too quickly or too slowly with respect to the total tree by using a custom R script; in this step, we omitted all trees with a relative branch length within the top and bottom 25% of this distribution (Figure S1). This step reduced the pool of global NFRs to 26,426 FASTA alignments in which the relative human branch length ranged between the lower and the upper quartiles.

To run our tests of selection, we fitted the null and alternative models for each DNA-element alignment using the batch scripts written by Haygood and collaborators (2007) that run under the program HyPhy (Pond et al. 2005). After all tests were completed, we extracted the best maximum likelihood estimates from twenty fittings to allow for stochasticity. Then, we obtained *P-*values from each likelihood ratio test (LRT) using the Chi^2^ distribution tool (*pchisq*) with one degree of freedom, implemented in R (R Team 2015). Interestingly, we observed that all *P*-value distributions were non-uniform and highly skewed to 1, therefore we considered our test to be conservative (Figure 3B). Consequently, we decided to use nominal *P*-values smaller than 0.05 to name regions scoring high for positive selection, instead of correcting for multiple testing to avoid violating the assumption of uniform distribution of the False Discovery Rate methodology (Benjamini and Hochberg 1995; Yekutieli and Benjamini 1999).

### Evaluation of the effect of query and reference length on sensitivity

To examine the sensitivity of our framework to detect positive selection under varying lengths of the query and reference, we obtained seven sets of 1000 queries of DHS alignments from the center of the peak position up to a total of 150 bp, 250 bp, 300 bp, 400bp, 500 bp, 600 bp and 1000 bp were reached in both sides. For each of these alignments, we also generated 10 reference alignments in order to account for the stochasticity of the evolutionary processes by concatenating 10 alignments from our set of 26K non-functional 300bp elements. Likewise, to identify the effects of variation in reference length in our queries, we also ran each query alignment of 300 bp against each of ten reference alignments varying from 300 bp to 30,000 bp. We also investigated the effects of including five species in our alignments rather than the minimum of three species considered by Haygood et al. (2007). There are now many genomewide assemblies that are available for many species, so we decided to employ additional species to test the effect of removing two branches in the tree. To do this, we extracted a random pool of 1,000 queries with both three (human, chimp and macaque) and five species (human, chimp, gorilla, orangutan and macaque) from the DHSs prepared by Thurman and collaborators (Thurman et al. 2012).

One typical concern of our method is the fact that it may be sensitive to deliver signals of selection if the reference sequences consist of negative selection, and if mutation rates are increased along all the branches of a tree. To test for these effects, we used the distribution of both relative and absolute branch lengths among non-functional sequences to simulate the distribution of trees under different classes of evolution (i.e. relaxation of constraint, increased mutation rate, positive selection or neutrality). To do this, we assumed that the alignments in the second quartile were putatively neutral, while those in the lower quartile were constrained, and those beyond the highest value of the upper quartile were under positive selection. Subsequently, we used these ‘neutral’ distributions to generate random sets of simulated trees, which were executed in the program *seq-gen* (Rambaut and Grassly 1997) to simulate sequence alignments in FASTA format. Consequently, we generated four sets of 1,000 query alignments in which 100% of the alignments were evolving by different classes of evolution: *i*, neutral in the foreground and in the background; *ii*, neutral in the foreground but constraint in the background by scaling down the substitution rates in the background by multiplying by a factor α = 0.1; and *iii*, scaling up the neutral rate in the foreground by a factor α = 5. Next, we used these simulated alignments to explore the power of our method to distinguish positive selection from relaxation of constraint and other types of evolution at the phylogenetic scale.

### Evaluation of sensitivity using known non-coding human accelerated elements

To test for selection in regions of the genome that have been previously investigated, we obtained a consolidated set of positive control queries of 2,649 non-coding human accelerated elements (ncHAEs) studied by Capra (2013). To compare the fraction of positive selection among ncHAEs with other sets of known regulatory elements, we used a subset of DHSs previously published by Thurman and collaborators (2012). To sample the DHSs in our test, we defined the “Ubiquity Score” as the fraction of cell types a given open chromatin site was open among the total number of tissues or cell types surveyed, thereby identifying a different set of queries to run our LRT method on known regulatory elements. To do this, we selected 4,216 DHSs that were open in at least 124 of 125 cell types, which we term ‘widespread sites’ (Ubiquity score > 0.98), and we selected 7,433 DHSs that were open in at most 2 of 125 cell types, which we term ‘specific sites’ (Ubiquity score < 0.02). Although these numbers seem arbitrary, they are a consequence of losing sequence alignments due to missing sequences from any of the branches or high frequency of gaps.

### Adapting the method to sliding windows

To test if our method is suitable for sliding windows, we applied *adaptiPhy* to test for positive selection within a 100 kb region around the NEIL3 locus, with the purpose of investigating additional signals of selection in the vicinity of a noncoding human accelerated element. Here, we split the entire region in windows of 300 bp with a 150 bp step size, and each query was run against a reference made from the filtered NFRs that were extracted from the masked genome alignment.

## DATA ACCESS

Sequence data presented in this manuscript is available in ENCODE. Scripts and software for analyzing data are available in Github (https://github.com/wodanaz/adaptiPhy). This GitHub repository contains a Docker file with the minimal requirements to run *adaptiPhy* in a sample dataset. Other datasets containing selection scores and neutral sequence regions are available in the supplementary data.

## Supporting information

Supplemental Images

## ACKNOWLEDGMENTS

A.B. was supported by a Hargitt Fellowship. We thank Dr. Andrew Allen and members in the Wray lab who provided feedback during the writing of this manuscript.

## AUTHOR CONTRIBUTIONS

A.B and G.A.W designed the study and wrote the manuscript. G.A.W and R. H. oversaw the project and analysis. A.B., R. H. and G.A.W analyzed and discussed. A.B., R. H., and G.A.W helped revise the manuscript.

## DISCLOSURE DECLARATION

All the authors declare no conflict of interest

## Notes

https://github.com/wodanaz/adaptiPhy

