## Supplemental Images for "Identifying branch-specific positive selection throughout the regulatory genome using an appropriate neutral proxy"

**SUPPLEMENTARY INFORMATION**

**Supplementary table 1.**

| **Locus** | **Nearest Gene** | **Distance from nearest gene** | **Function of the nearest gene** |
| --- | --- | --- | --- |
| chr2:236773664-236774209 | AGAP1 | 370.9 kb | GTP binding and phospholipid binding and endocytosis |
| chr4:178272899-178273376 | NEIL3 | 41.9 kb | Nucleic acid binding and single-stranded DNA binding and telomere C-strand synthesis. |
| chr9:16717947-16718304 | BNC2 | 307.7 kb | Zinc finger protein functioning in skin color saturation and associated with idiopathic scoliosis. |
| chr16:82295069-82295368 | MPHOSPH6 | 91.5 kb | Deadenylation-dependent mRNA decay and rRNA processing in the nucleus and cytosol. |

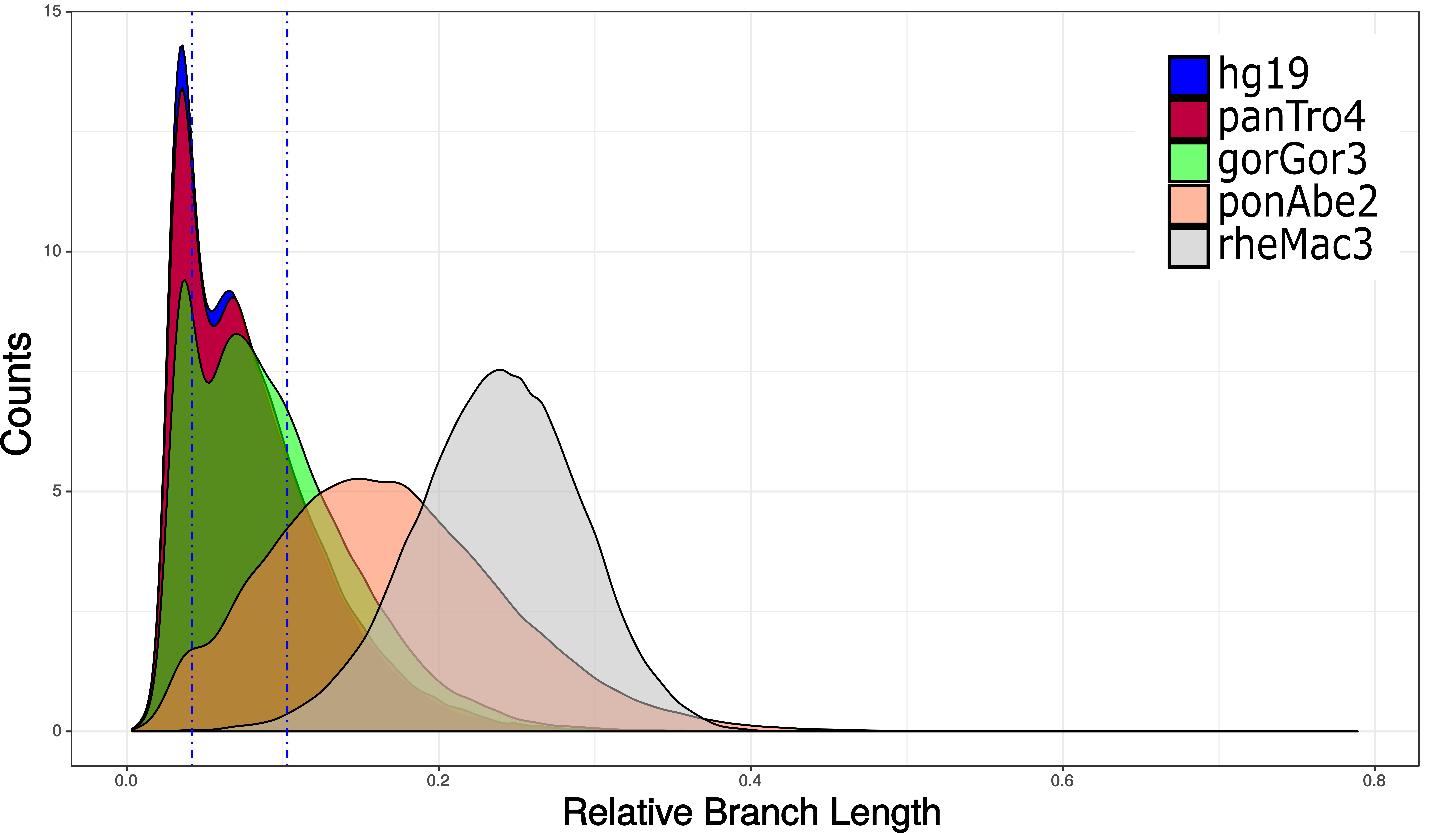

**Fig S1:** Distribution of relative branch length across non-functional and putatively neutrally evolving elements.

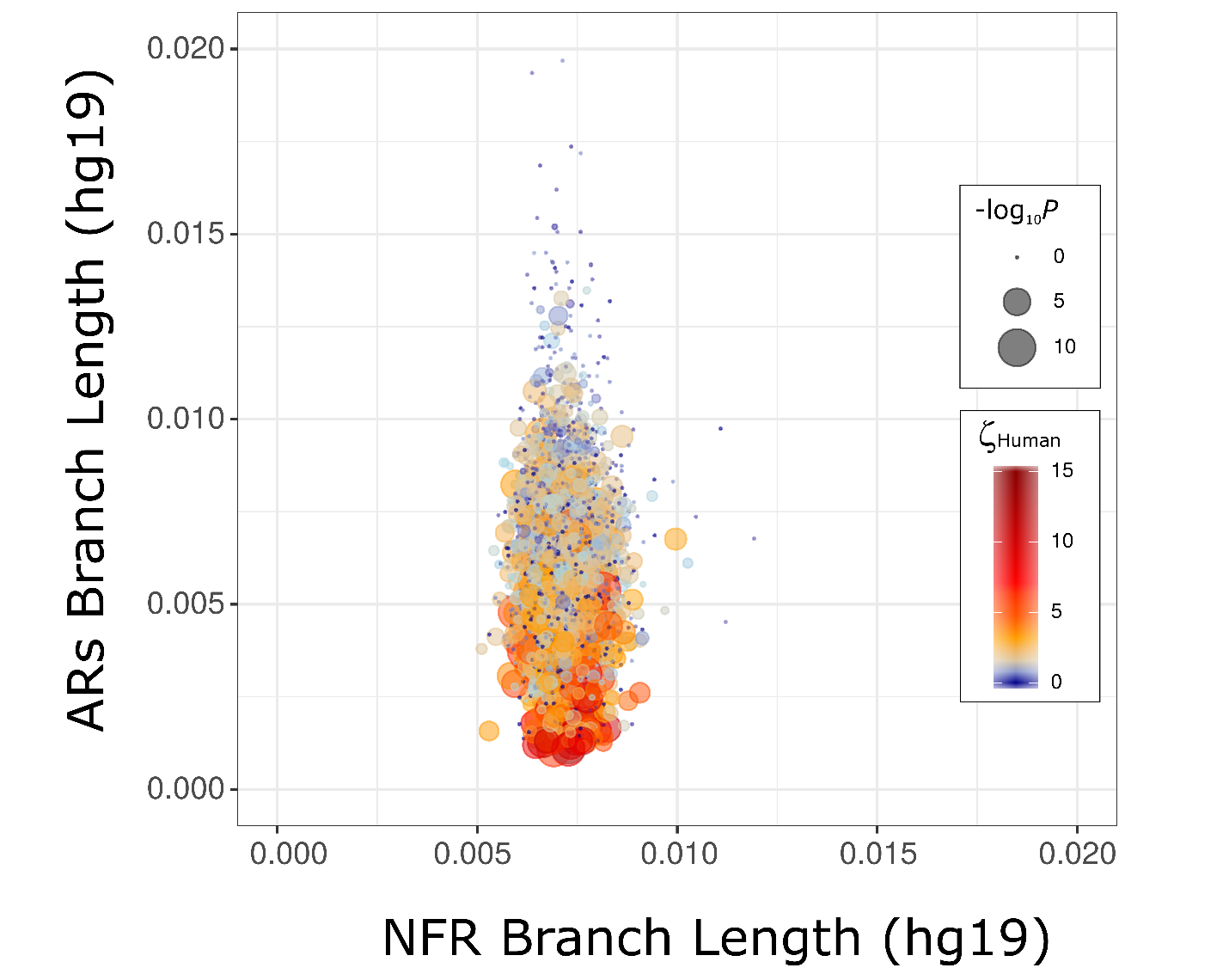

**Fig S2. AR elements bias the estimation of positive selection.** The distribution of substitution rates among ARs (y-axis) is appreciably wider than in post-filtered non-functional regions (x-axis). This increases the rate of false positives in AR regions that are more conserved. Color gradient depicts the magnitude of ζ while size depicts the significance (-Log_10_ *P*value). Cold colors depicts regions that are neutral or constrained, while warmer colors depict more hits of positive selection.

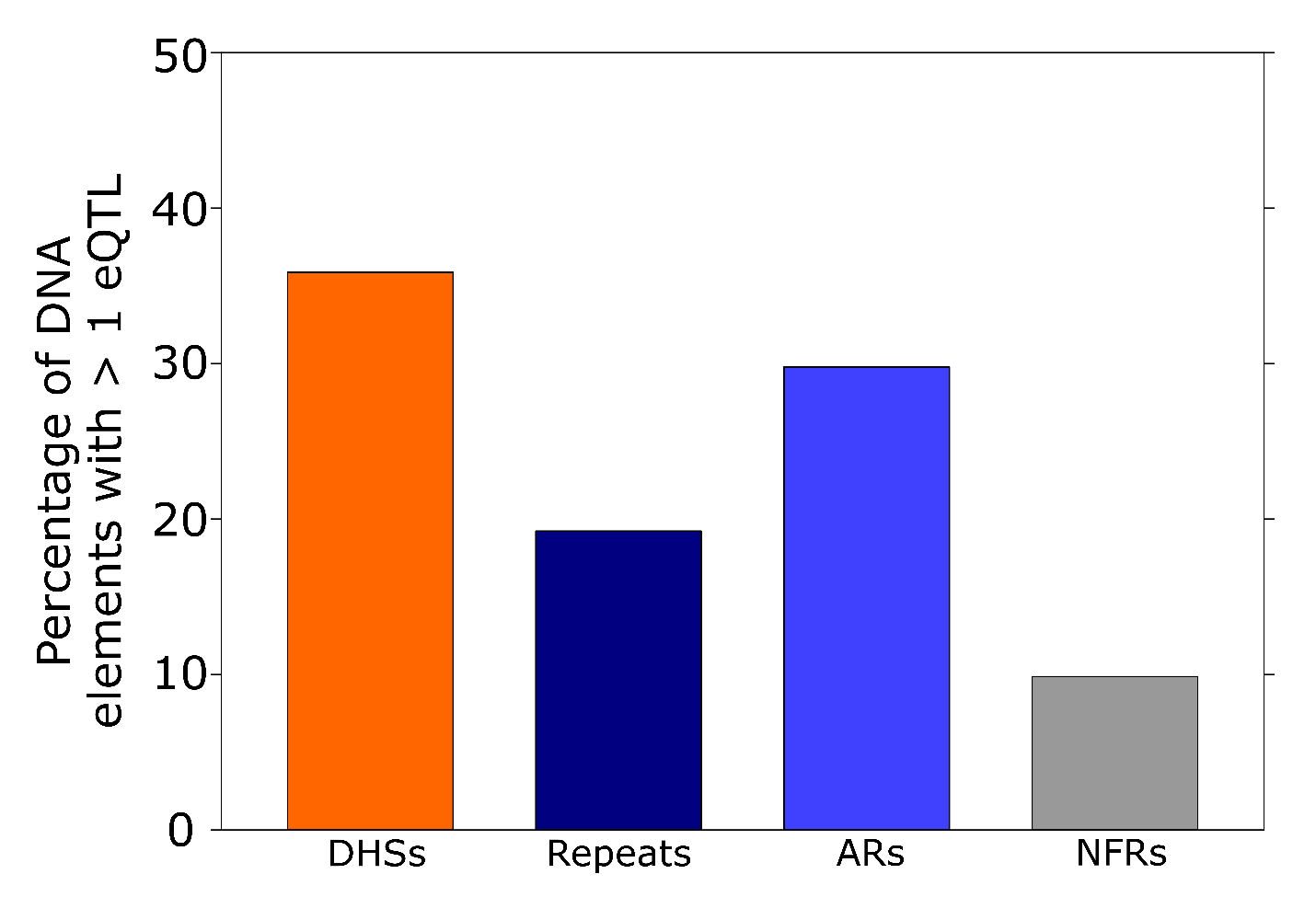

**Fig S3. Percentage of DNA elements with at least one multi-tissue eQTL.** Many repeats and ancestral repeats contain at least 1 multitissue eQTL. Almost 9% of our random list of 5104 non-functional elements contain at least 1 eQTL.

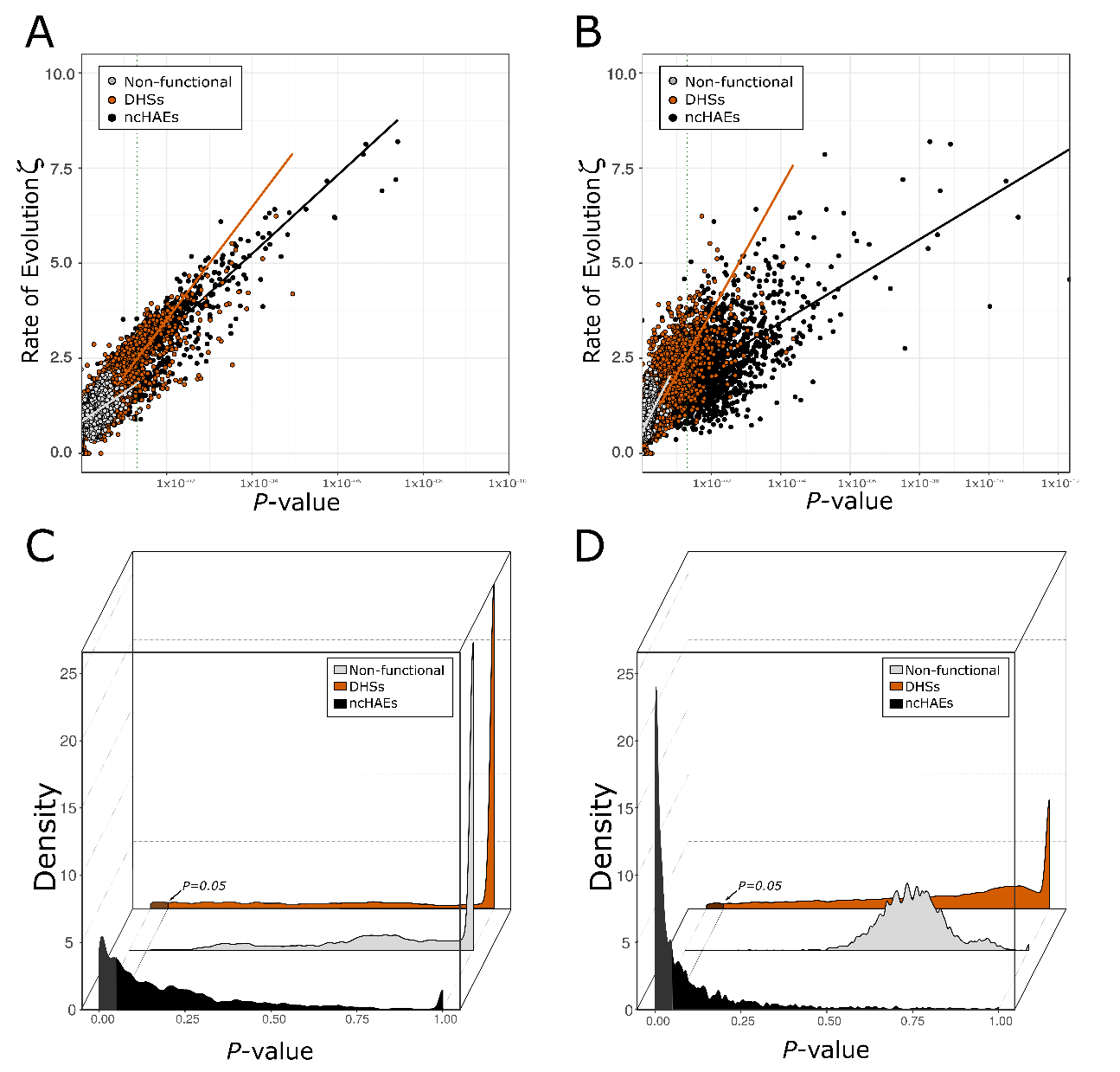

**Fig S4: Correlation between evolutionary ratio and *P*-value.** Rate of evolution in the human branch using global neutral proxy for DHSs (orange), non-functional elements (gray), and noncoding ncHAEs (black). The green dotted line depicts significance at the 0.05 level. **A.** All the Spearman correlation coefficients for our framework using *HyPhy* are strongly significant (*P* < 2.2 x10^-16^), and very high (rho = 0.86 for ncHAEs and rho = 0.80 for DHSs) except for non-functional data (rho = 0.60). **B.** All the Spearman correlation coefficients for *phyloP* scores of acceleration using our global neutral proxy are also strongly significant (*P* < 2.2 x10^-16^), and very high (rho = 0.55 for ncHAEs and rho = 0.90 for DHSs) except for non-functional data (rho = 0.47). **C-D.** Density distributions of *P*-value among different classes of DNA elements including non-functional sequences (gray); a set of 11649 DHSs (orange) from Thurman et al (2012); and the distribution of *P*-values provided by our tests for ncHAEs (black) for *adaptiPhy* **(C)** and *phyloP* scores of acceleration **(D)**. Significant distributions of elements scoring high for positive selection have been highlighted.

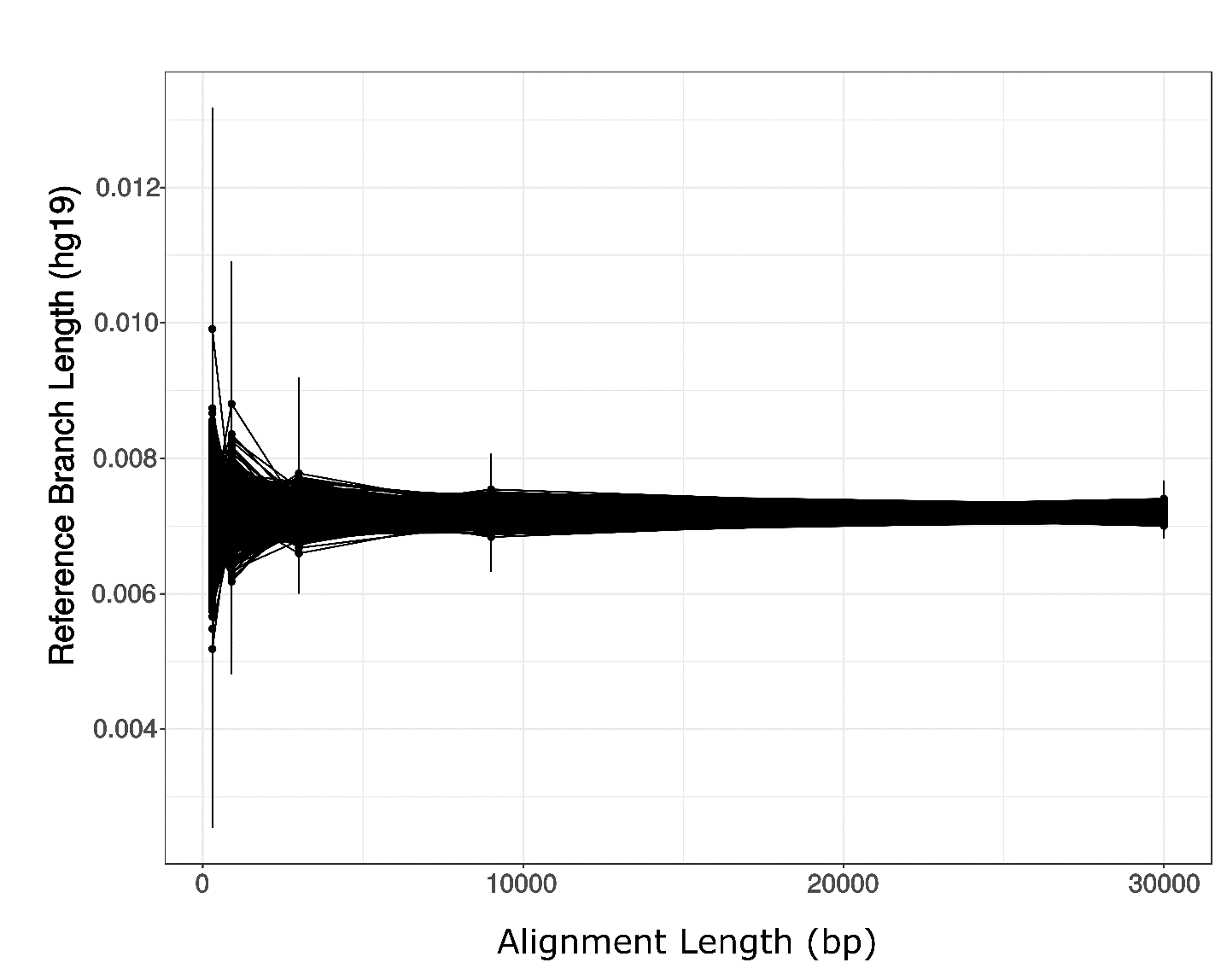

**Fig S5. Effect of alignment length on reference branch length.** Variation of branch length in the human branch relative to the alignment length among concatenated references from real non-coding data (black).

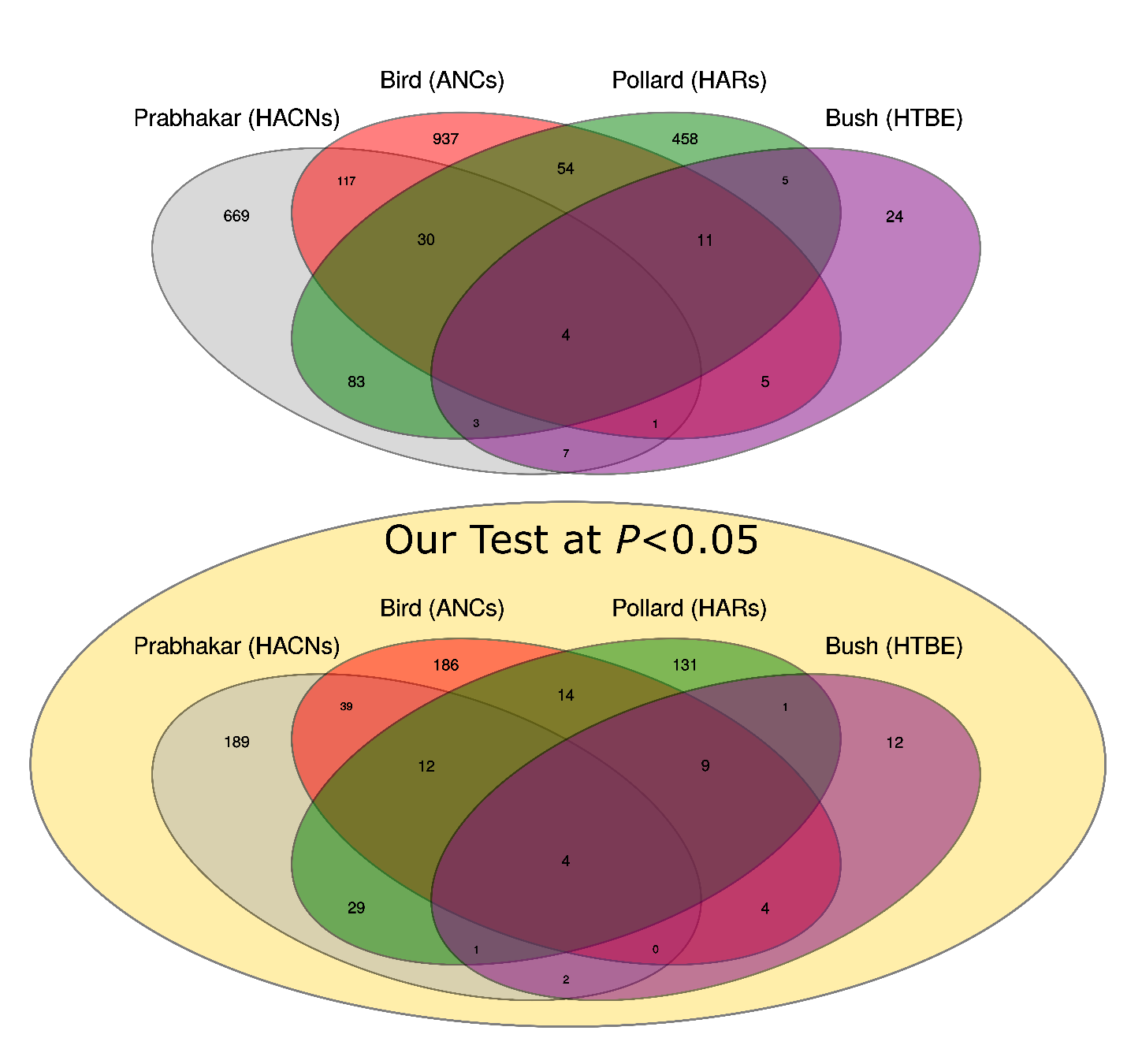

**Fig S6: Overlaps among different datasets of human accelerated elements.** **A.** Total overlap between HACNs, ANCs, HARs and HTBEs (Pollard, Salama, King, et al. 2006; Bird et al. 2007; Bush and Lahn 2008; Prabhakar et al. 2008). **B.** Overlap among all human accelerated elements in A that scored high for positive selection with *adaptiPhy* at a *P* < 0.05.
